## Supplementary Information for "Marine heatwaves disrupt ecosystem structure and function via altered food webs and energy flux"

**The PDF file includes:**

Figures S1 to S5

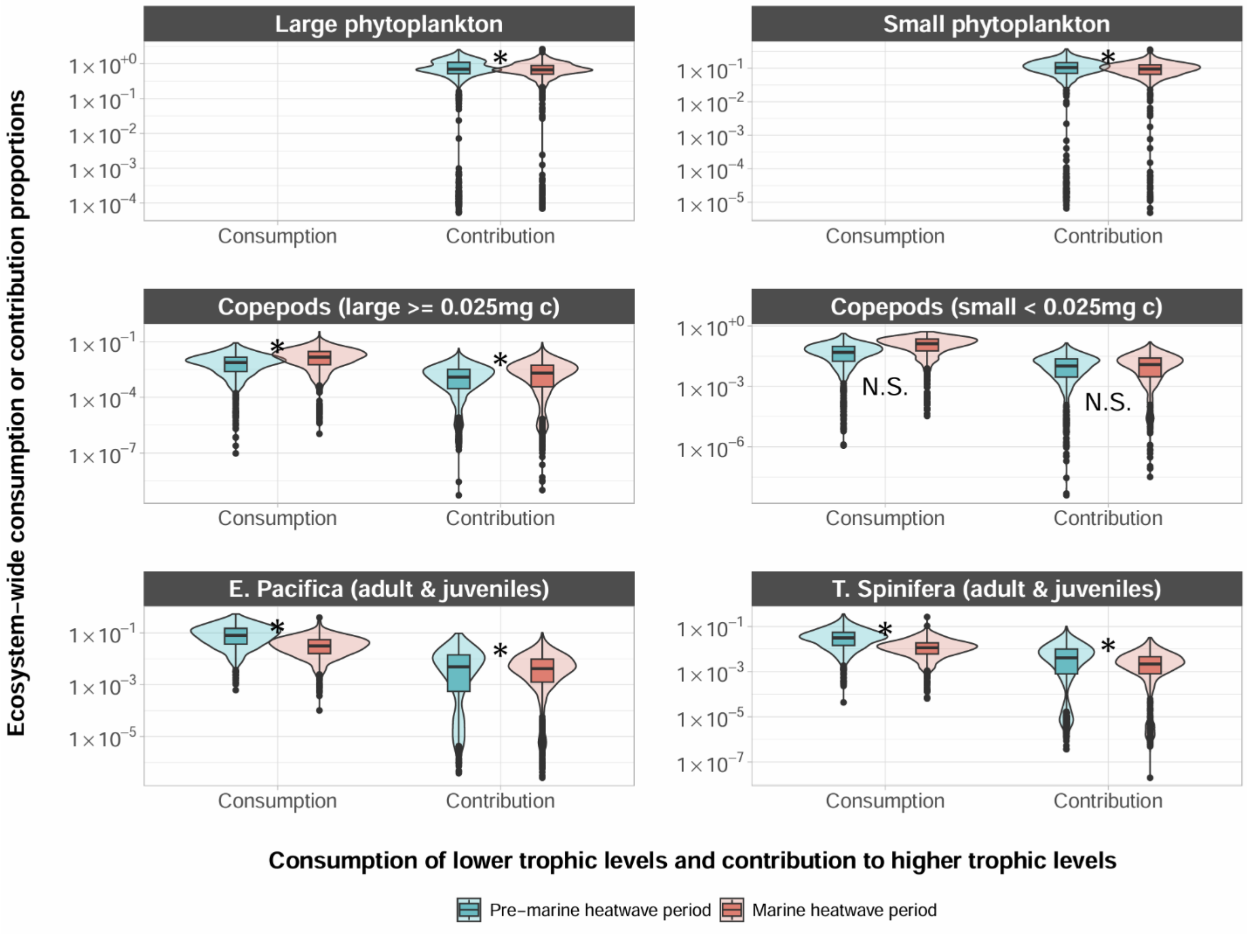


**Figure S1. Ecosystem-wide consumption of prey and contribution to predators of lower trophic levels.**

Violin plots show density of points from 1000 Monte Carlo runs of ecosystem models. Plotted over the violin plots, boxplots show median values as thick horizontal lines and first and third quartiles (the 25th and 75th percentiles) as the lower and upper edges of the box, respectively (n = 1000 independent Monte Carlo model parameterizations; see Methods). The lower and upper whiskers extend from the edges of the box to the values that are smallest and largest (respectively), yet no further than 1.5 × interquartile range (i.e., the distance between the first and third quartiles) from the box. Outlying data beyond the end of the whiskers are plotted as individual points. Asterisks indicate that the difference in consumption of prey and contribution to predators between the pre-MHW and MHW models is significantly different (exact p-values found in Source Data file). Statistical significance was determined via t-tests with Bonferroni corrections for multiple comparisons. Units are proportions of total ecosystem consumption or contributions (see notes on footprint and reach in the Methods). Source data are provided as a Source Data file.

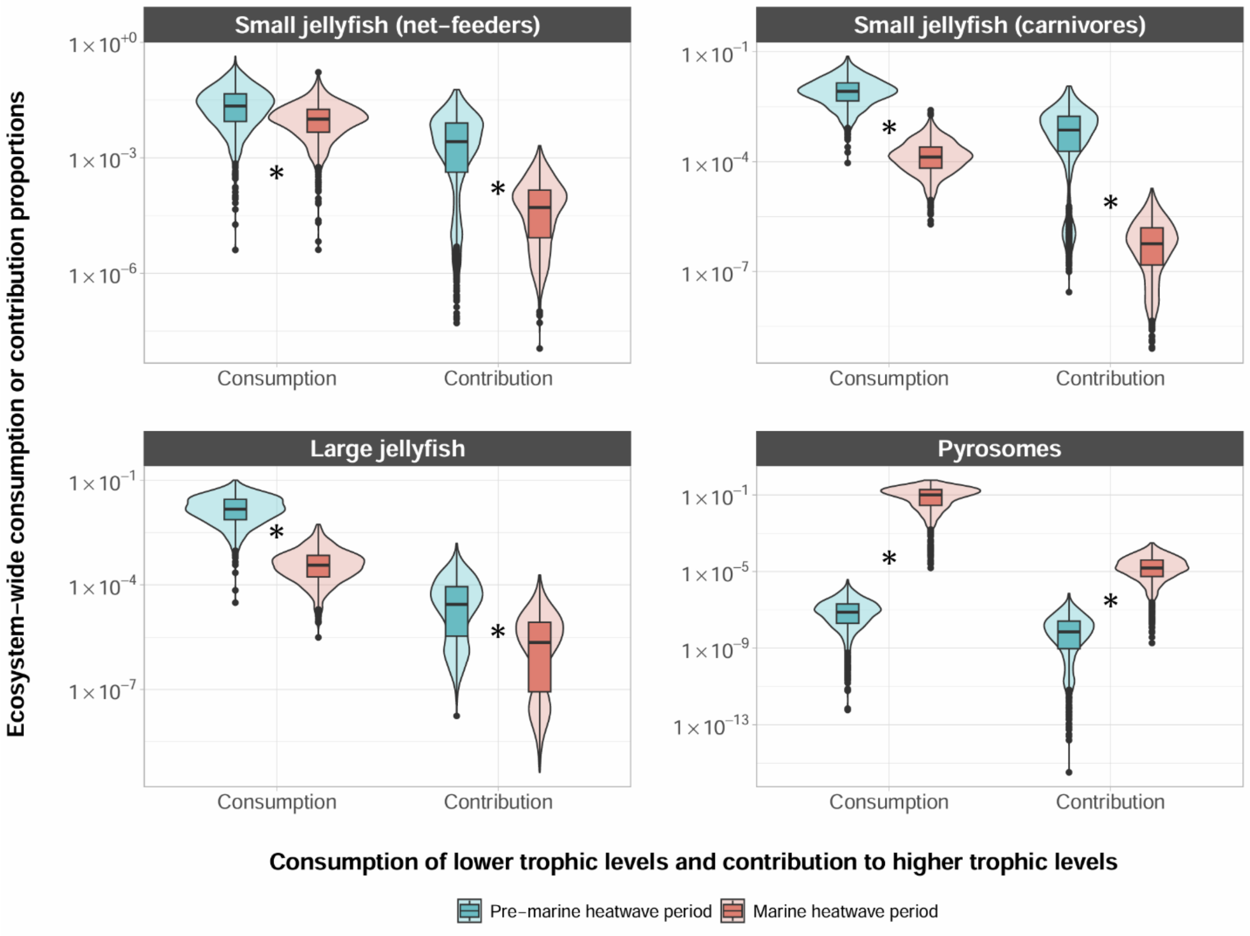


**Figure S2. Ecosystem-wide consumption of prey and contribution to predators of gelatinous animals.**

Violin plots show density of points from 1000 Monte Carlo runs of ecosystem models. Plotted over the violin plots, boxplots show median values as thick horizontal lines and first and third quartiles (the 25th and 75th percentiles) as the lower and upper edges of the box, respectively (n = 1000 independent Monte Carlo model parameterizations; see Methods). The lower and upper whiskers extend from the edges of the box to the values that are smallest and largest (respectively), yet no further than 1.5 × interquartile range (i.e., the distance between the first and third quartiles) from the box. Outlying data beyond the end of the whiskers are plotted as individual points. Asterisks indicate that the difference in consumption of prey and contribution to predators between the pre-MHW and MHW models is significantly different (exact p-values found in Source Data file). Statistical significance was determined via t-tests with Bonferroni corrections for multiple comparisons. Units are proportions of total ecosystem consumption or contributions (see notes on footprint and reach in the Methods). Source data are provided as a Source Data file.

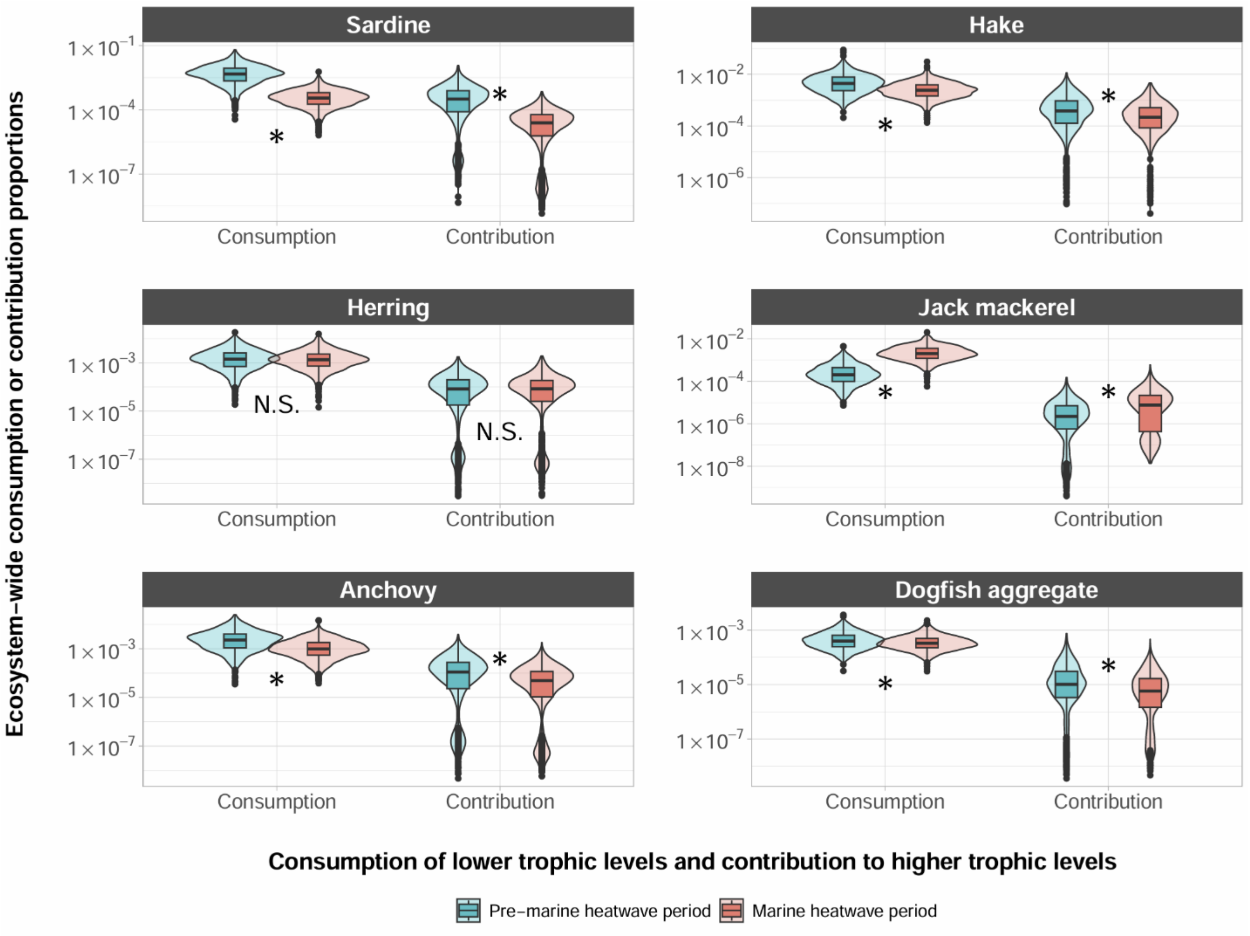


**Figure S3. Ecosystem-wide consumption of prey and contribution to predators of various fishes.**

Violin plots show density of points from 1000 Monte Carlo runs of ecosystem models. Plotted over the violin plots, boxplots show median values as thick horizontal lines and first and third quartiles (the 25th and 75th percentiles) as the lower and upper edges of the box, respectively (n = 1000 independent Monte Carlo model parameterizations; see Methods). The lower and upper whiskers extend from the edges of the box to the values that are smallest and largest (respectively), yet no further than 1.5 × interquartile range (i.e., the distance between the first and third quartiles) from the box. Outlying data beyond the end of the whiskers are plotted as individual points. Asterisks indicate that the difference in consumption of prey and contribution to predators between the pre-MHW and MHW models is significantly different (exact p-values found in Source Data file). Statistical significance was determined via t-tests with Bonferroni corrections for multiple comparisons. Units are proportions of total ecosystem consumption or contributions (see notes on footprint and reach in the Methods). Source data are provided as a Source Data file.

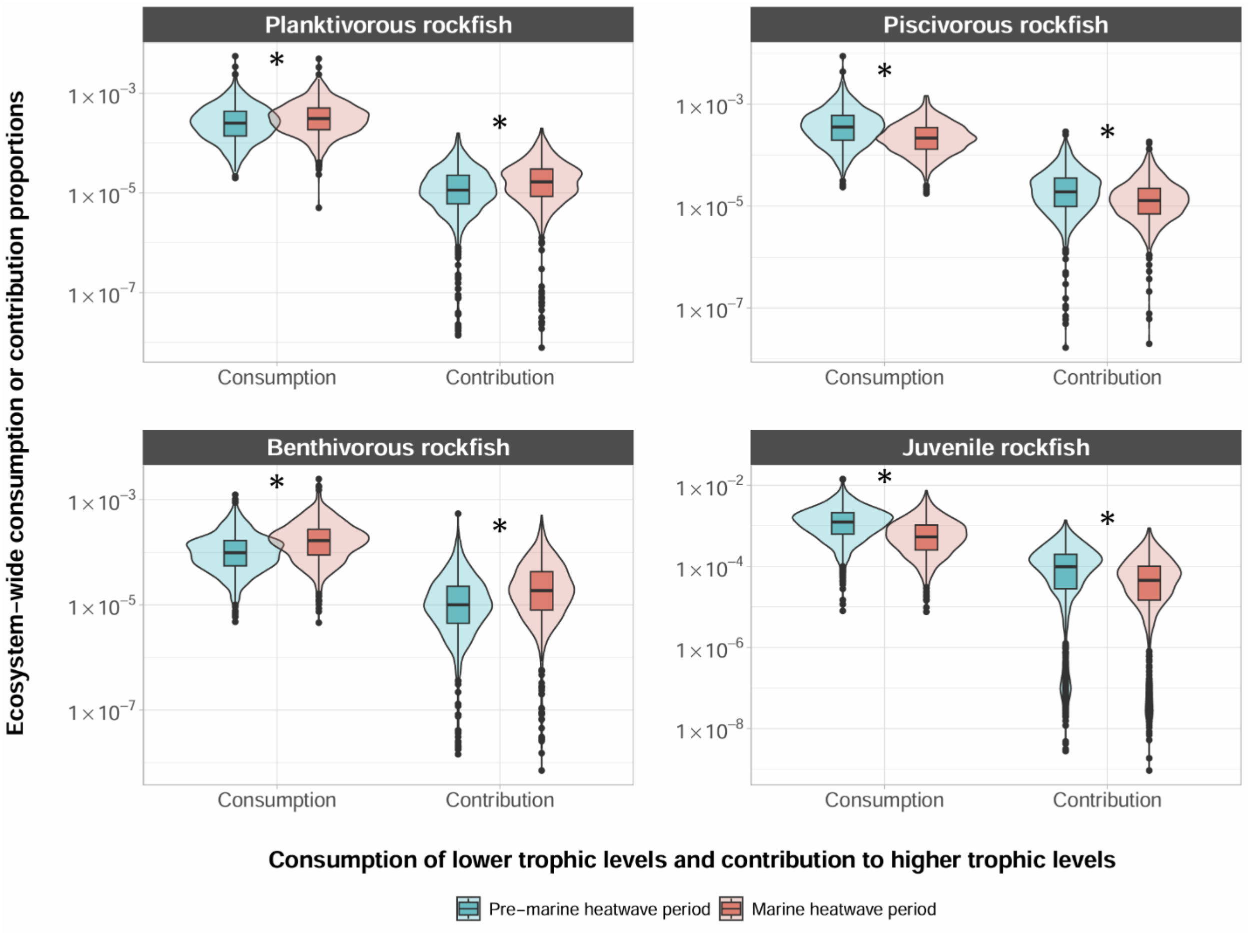


**Figure S4. Ecosystem-wide consumption of prey and contribution to predators of rockfishes.**

Violin plots show density of points from 1000 Monte Carlo runs of ecosystem models. Plotted over the violin plots, boxplots show median values as thick horizontal lines and first and third quartiles (the 25th and 75th percentiles) as the lower and upper edges of the box, respectively (n = 1000 independent Monte Carlo model parameterizations; see Methods). The lower and upper whiskers extend from the edges of the box to the values that are smallest and largest (respectively), yet no further than 1.5 × interquartile range (i.e., the distance between the first and third quartiles) from the box. Outlying data beyond the end of the whiskers are plotted as individual points. Asterisks indicate that the difference in consumption of prey and contribution to predators between the pre-MHW and MHW models is significantly different (exact p-values found in Source Data file). Statistical significance was determined via t-tests with Bonferroni corrections for multiple comparisons. Units are proportions of total ecosystem consumption or contributions (see notes on footprint and reach in the Methods). Source data are provided as a Source Data file.

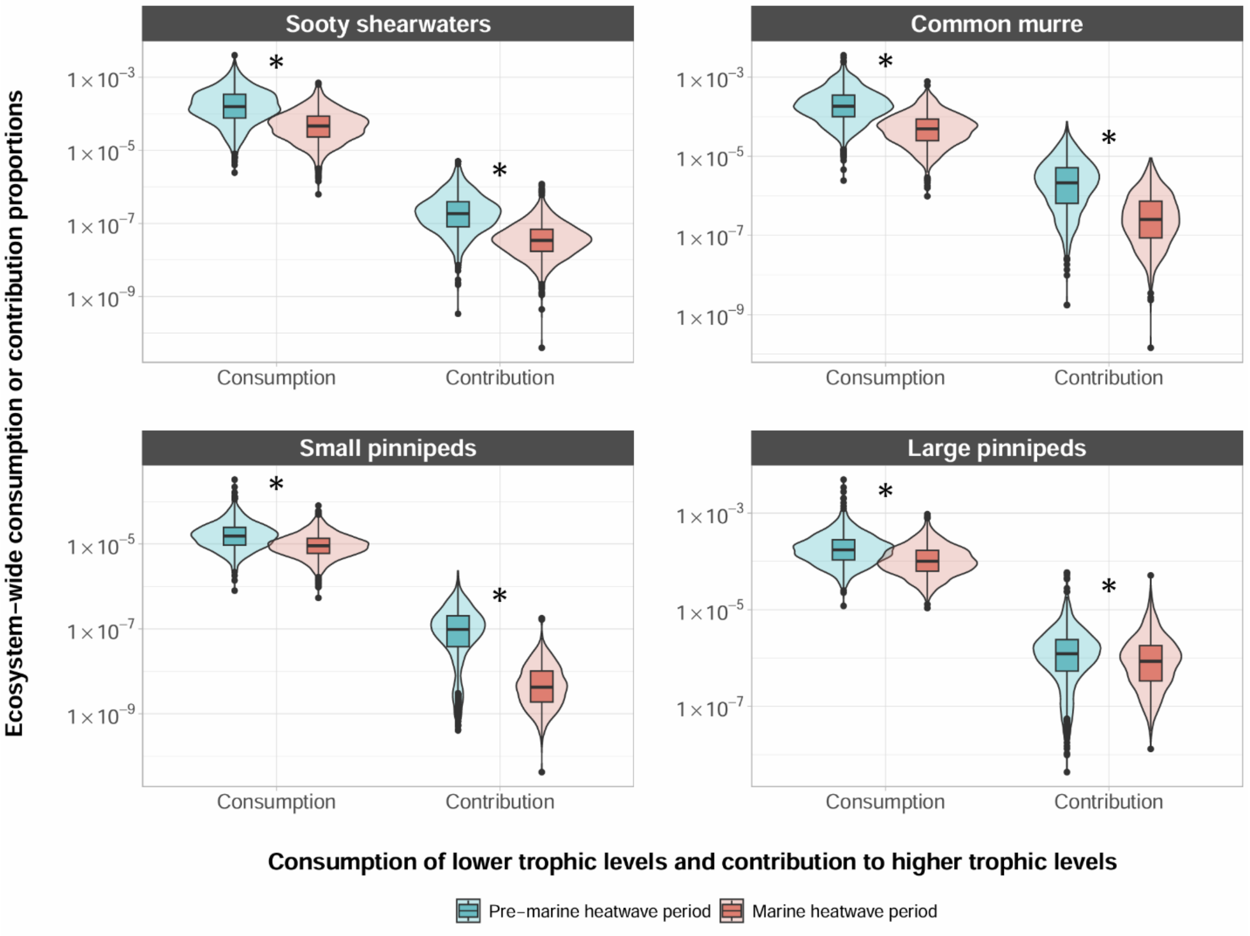


**Figure S5. Ecosystem-wide consumption of prey and contribution to predators of seabird and mammal predators.**

Violin plots show density of points from 1000 Monte Carlo runs of ecosystem models. Plotted over the violin plots, boxplots show median values as thick horizontal lines and first and third quartiles (the 25th and 75th percentiles) as the lower and upper edges of the box, respectively (n = 1000 independent Monte Carlo model parameterizations; see Methods). The lower and upper whiskers extend from the edges of the box to the values that are smallest and largest (respectively), yet no further than 1.5 × interquartile range (i.e., the distance between the first and third quartiles) from the box. Outlying data beyond the end of the whiskers are plotted as individual points. Asterisks indicate that the difference in consumption of prey and contribution to predators between the pre-MHW and MHW models is significantly different (exact p-values found in Source Data file). Statistical significance was determined via t-tests with Bonferroni corrections for multiple comparisons. Units are proportions of total ecosystem consumption or contributions (see notes on footprint and reach in the Methods). Source data are provided as a Source Data file.
